# Downstream effects of the “Less, but More” Fgf signaling in *Oikopleura dioica*: Fgf receptor expansion and RTK pathway simplification

**DOI:** 10.64898/2025.12.09.693154

**Authors:** Gaspar Sánchez-Serna, Paula Bujosa, Alfonso Ferrández-Roldán, Marc Fabregá-Torrus, Ana Alonso-Bartolomé, Laura Reyner-Laplana, Nuria P. Torres-Águila, Cristian Cañestro

**Author notes:** corresponding author: Cristian Cañestro < >. Gaspar Sánchez-Serna < >, Paula Bujosa < >, Alfonso Ferrández-Roldán < >, Ana Alonso Bartolomé < >, Laura Reyner Laplana < >, Marc Fabrega-Torrus < >, Nuria P. Torres-Águila < >.

## Abstract

Fibroblast growth factor (FGF) signaling is central to chordate development and has been extensively remodeled in tunicates. Recent findings show that appendicularians have massively lost all ancestral chordate Fgf subfamilies except two, the Fgf9/16/20 and Fgf11/12/13/14 subfamilies, which in contrast have undergone a burst of lineage-specific duplications and diversification into novel paralogs, in an evolutionary scenario that we have named “Less, but More”. Here, we investigate the downstream effects of the Fgf losses and duplications ion Fgf receptors (FgfRs) and intracellular RTK components in the appendicularian *Oikopleura dioica*. We show that the single ancestral FgfR gene has expanded into three paralogs (FgfRa–c), which are conserved across cryptic *O. dioica* species, yet highly divergent from other chordates. Despite strong sequence divergence, structural modeling indicates preservation of canonical FgfR architecture. Expression analyses reveal distinct spatiotemporal patterns: *FgfRa* and *FgfRb* are maternally supplied and enriched in mesodermal derivatives, whereas FgfRc is restricted to neural and epithelial tissues. Genome surveys of downstream RTK pathways show conservation of core RAS/MAPK, PLCγ/PKC, and PI3K/AKT cascades, but with losses of classical Ras genes and several adaptors, suggesting a lineage-specific simplification of transduction complexes. Transduction gene expression shifts from broad maternal ubiquity to tissue-specific domains, particularly in brain, notochord, muscle, and gonadal primordia throughout embryonic and larval development. Appendicularians appear as the only non-vertebrate chordate lineage that recapitulate the vertebrate-like FgfR expansion following Fgf ligands diversification. Downstream components, however, evolved more conservatively, tending toward simplification, reinforcing the view that appendicularians generate signaling innovation despite extensive gene loss.

## INTRODUCTION

Fibroblast growth factors (FGF) and their receptors (FgfR) constitute a deeply conserved signaling pathway involved in numerous biological processes in eumetazoans, including embryonic development, morphogenesis, and cell fate decisions (Teven et al. 2014; Itoh et al. 2016). The binding of secreted Fgfs to FgfRs on the cell surface activates one or several intracellular transduction pathways, being the best-known the PI3K, PLCy, and RAS/MAPK pathways (Goetz and Mohammadi 2013). Understanding the evolution of the FGF signaling pathway is challenging because both its components and its functions vary widely across animal groups. However, its presence in all eumetazoans and its consistent involvement in key developmental processes, together with its recruitment into many derived and lineage-specific roles, make it an especially interesting system for studying how the evolution of animal body plans is linked to the diversification of their underlying genetic toolkits (Oulion et al. 2012; Bertrand et al. 2014).

Within the chordate phylum, the ancestral chordate possessed a single FgfR gene and at least eight Fgf genes, one representative of each of the current subfamilies described in vertebrates. While the vertebrate lineage expanded the Fgf and FgfR repertoire through two rounds of whole-genome duplication - a process that has been associated to the evolutionary acquisition of vertebrate-specific innovations-, cephalochordates and ascidian tunicates retained a basal complement similar to that of the ancestral chordate (Satou et al. 2002; Bertrand et al. 2011; Oulion et al. 2012). Our previous work demonstrated that appendicularian tunicates have massively lost all ancestral chordate Fgf subfamilies except two. This reduction was accompanied by a burst of duplications that expand and diversify the remaining Fgf9/16/20 and Fgf11/12/13/14 subfamilies, resulting in novel paralogs with distinct developmental functions (Sánchez-Serna et al. 2025). On this basis, we proposed the “Less, but more” scenario, which characterizes evolutionary contexts in which extensive gene loss co-occurs with the expansion and functional diversification of the surviving genes. The combined action of these processes can give rise to four outcomes: retention of ancestral functions, elimination of functions associated with lost genes, redistribution of functions between lost and novel genes, and the emergence of novel functions. This scenario emphasizes that gene loss, in conjunction with subfunctionalization and neofunctionalization of duplicates associated to the losses, can constitute a major force in the evolution and diversification of biological lineages.

Appendicularians represent the second major tunicate lineage alongside ascidians and thaliaceans. Unlike their relatives, appendicularians retain a largely larva-like body plan throughout their life cycle. They are small, free-swimming planktonic organisms that inhabit pelagic marine environments and are characterized by the construction of complex extracellular “houses” used for filter feeding. Despite their morphological simplicity, appendicularians exhibit some of the most accelerated genomic evolution known among chordates. *Oikopleura dioica*, the best-studied representative, possesses a compact genome with extensive gene loss, high sequence divergence, and numerous genomic rearrangements, while nonetheless maintaining a conserved chordate body plan and core developmental processes (Nishida 2008; Denoeud et al. 2010; Ferrández-Roldán et al. 2019; Plessy et al. 2024). This combination of fast-evolving genomes and conserved morphology makes appendicularians a powerful model for investigating how developmental systems adapt to large-scale modifications in gene content (Ferrández-Roldán et. 2021). Their unusual genomic architecture, together with their phylogenetic position as close relatives of vertebrates, provides a unique opportunity to examine how the FGF signaling pathway, and more broadly, gene regulatory and signaling networks have been remodeled during chordate evolution.

Our findings on the evolution of the Fgf gene family directed the focus of our attention to the evolution of the remaining components of the Fgf signaling pathway in appendicularians. These are the Fgf receptor (FgfR) and the downstream components related to the transduction of the extracellular signal inside the cells. Here, we characterize the evolution, structural features, and expression dynamics of FgfR paralogs in *O. dioica*, and assess the conservation and developmental expression of the three canonical FgfR transduction cascades: RAS/MAPK, PLCγ/PKC, and PI3K/AKT.

## MATERIALS AND METHODS

### Laboratory culture of *Oikopleura dioica*

*O. dioica* specimens were obtained from laboratory colonies maintained at the University of Barcelona for over five years. The original founder individuals were collected from the Mediterranean coast near Barcelona (Catalonia, Spain) and cultured following the protocols described in Martí-Solans et al. (2015). This work did not raise ethical concerns, as research on aquatic invertebrates is not subject to animal experimentation regulations under Real Decreto 223/1998 or Catalonia Ley 5/1995, DOGC 2073, 5172. Nevertheless, all procedures complied with European Union (EU) guidelines for animal care and were formally approved by the Ethical Animal Experimentation Committee (CEEA-2009) of the University of Barcelona.

### Genome database searches, gene identification and phylogenetic analyses

FgfR genes in *O. dioica* were initially detected in the reference genome database corresponding to the Norwegian population (Danks et al. 2013), using BLASTp and tBLASTn searches, with *Homo sapiens* and *Ciona robusta* FgfR protein sequences as queries. These candidate genes were subsequently located using tBLASTn in the genomic assemblies of the Barcelona (BAR), Osaka (OSA), and Okinawa (OKI) *O. dioica* lineages (Plessy et al. 2024). Within each lineage, the identified FgfR sequences were then used as queries to search for additional paralogs, allowing us to compile the final FgfR gene set for each genome.

To characterize FgfR genes in other appendicularian species, we conducted tBLASTn searches using the *O. dioica* FgfR proteins against publicly available genomes (*Oikopleura albicans*, *Oikopleura vanhoeffeni*, *Oikopleura longicauda*, *Bathochordaeus sp*., and *Mesochordaeus erythrocephalus*) (Naville et al. 2019). For ascidian species beyond *C. robusta*, FgfR sequences from *C. robusta* were used as BLASTp and tBLASTn queries against gene models and genome assemblies accessible through ANISEED, including *Ciona savignyi*, *Phallusia fumigata, Phallusia mammillata, Halocynthia roretzi, Halocynthia aurantium, Botryllus schlosseri, Botryllus leachii, Molgula occulta, Molgula oculata*, and *Molgula occidentalis* (Brozovic et al. 2018).

Alignments of the conserved tyrosine kinase domain were produced with MUSCLE and MAFFT using Aliview v1.28 (Larsson 2014), and manually inspected. Phylogenetic relationships were inferred using Maximum Likelihood (ML) methods implemented in PhyML v3.0 (Guindon et al. 2010), and IQ-Tree (Nguyen et al. 2015). Node support was assessed with the SH-like aLRT method and ultrafast bootstrap replicates (n = 100).

### Protein structure analyses

The domain composition and functional motifs of FgfR proteins were analyzed individually using InterProScan to obtain an in silico functional annotation of each sequence (Jones et al. 2014). Pairwise sequence identity and similarity were calculated using EMBOSS Needle based on global alignments (Madeira et al. 2024). Three-dimensional structures of *O. dioica* FgfR proteins were generated de novo with AlphaFold2 (Jumper et al. 2021). For each receptor, the highest-ranked relaxed model was imported into UCSF ChimeraX for visualization, structural inspection, and figure preparation (Pettersen et al. 2021).

### Cloning and expression analyses

*O. dioica* FgfR and transduction pathway genes were PCR-amplified from cDNA or gDNA extracted from individuals of the Barcelona population, following the procedures described in Martí-Solans et al. (2016). PCR products were cloned using the TOPO TA Cloning Kit (K4530-20, Invitrogen), and the resulting plasmids were digested with the appropriate restriction enzymes to generate antisense DIG-labeled riboprobes for whole-mount in situ hybridization (WMISH) (Bassham and Postlethwait 2000; Cañestro and Postlethwait 2007; Martí-Solans et al. 2016).

Details on probe sequences, primer sets, templates, product lengths, and RNA polymerases/restriction enzymes used are provided in Supplementary Table 1.

## RESULTS

### Evolution of the Fgf receptor in appendicularians

Genomic searches by RBBH and phylogenetic analyses allowed us to identify three *FgfR* genes in *Oikopleura dioica* and several orthologs in all other appendicularian species surveyed (**Figure 1**). Considering that ascidians and cephalochordates each had a single *FgfR* gene (Kamei et al. 2000; Satou et al. 2003; D’Aniello et al. 2008), and that the four vertebrate *FgfR* paralogs resulted from two rounds of whole genome duplication specific to the vertebrate lineage (Itoh and Ornitz 2004), the three *O. dioica FgfR* genes appeared as paralogs resulting from a lineage-specific expansion in appendicularians after their split from the rest of tunicates.

**Figure 1.**
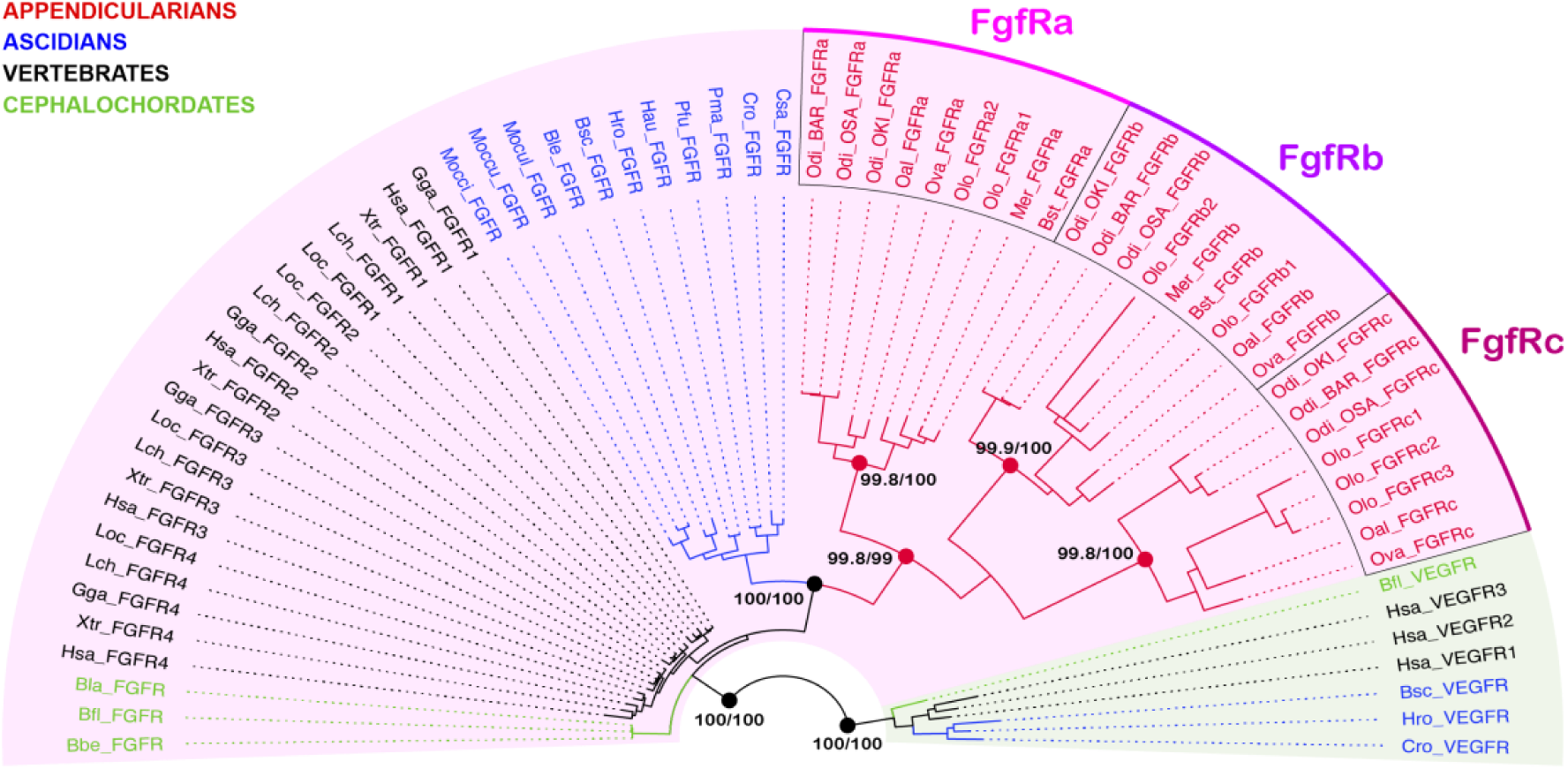
Evolutionary relation of appendicularian Fgf receptors. Maximum likelihood phylogenetic tree of chordate FgfR TK domains (shaded in magenta) including the orthologs found in appendicularians. Cephalochordate orthologs are depicted in green, vertebrate orthologs in black, ascidian orthologs in blue, and appendicularian orthologs in red. VEGFR TK domains were added as an outgroup, shaded in green. Node support values correspond to SH-aLRT support (%) / ultrafast bootstrap support (%).

The tree topology suggested that the ancestral appendicularian *FgfR* gene (*FgfRabc*) was duplicated into two paralogs (*FgfRa* and *FgfRbc*), and subsequently one of these paralogs (*FgfRbc*) underwent a second duplication in the *Oikopleurinae* lineage (i.e. *O. dioica*, *O. albicans* and *O. vanhoeffeni*), while in the *Bathochordaeinae* lineage the presence of a single FgfRbc gene in *Bathochordaeus spp*. and *Mesochordaeus spp*., but multiple copies in *O. longicauda*, hindered the evolutionary reconstruction. Species-specific *FgfR* duplications in *O. longicauda* indicated that in some appendicularians the *FgfR* catalogue might be composed of up to at least seven genes. This suggested that the *FgfR* gene in appendicularians has undergone a process of expansion and diversification in parallel to that of the only two surviving Fgf9/16/20 and Fgf11/12/13/14 subfamilies (Sánchez-Serna et al. 2025).

### Conservation and divergence in appendicularians FgfR genes

In *O. dioica*, the three FgfR paralogs have undergone extensive sequence divergence, both relative to each other and to other chordate FgfR proteins. Nevertheless, *in silico* protein modeling revealed that, despite this divergence, all three *O. dioica* FgfRs retain structural features characteristic of canonical FgfRs. Each receptor comprised two extracellular immunoglobulin-like (IG-like) domains, a transmembrane helix, and an intracellular split tyrosine kinase (TK) domain (**Figure 2**). This two–IG-like domain configuration contrasted with the three IG-like domains typically found in vertebrate and cephalochordate FgfRs (D’Aniello et al. 2008; Ornitz and Itoh 2015), but mirrored what has been reported for *Ciona robusta* and other ascidian FgfRs (Shimauchi et al. 2001; Satou et al. 2003). These observations indicated that the loss of IG-like domain 1 (IG-L1) was an ancestral event in the tunicate lineage.

**Figure 2.**
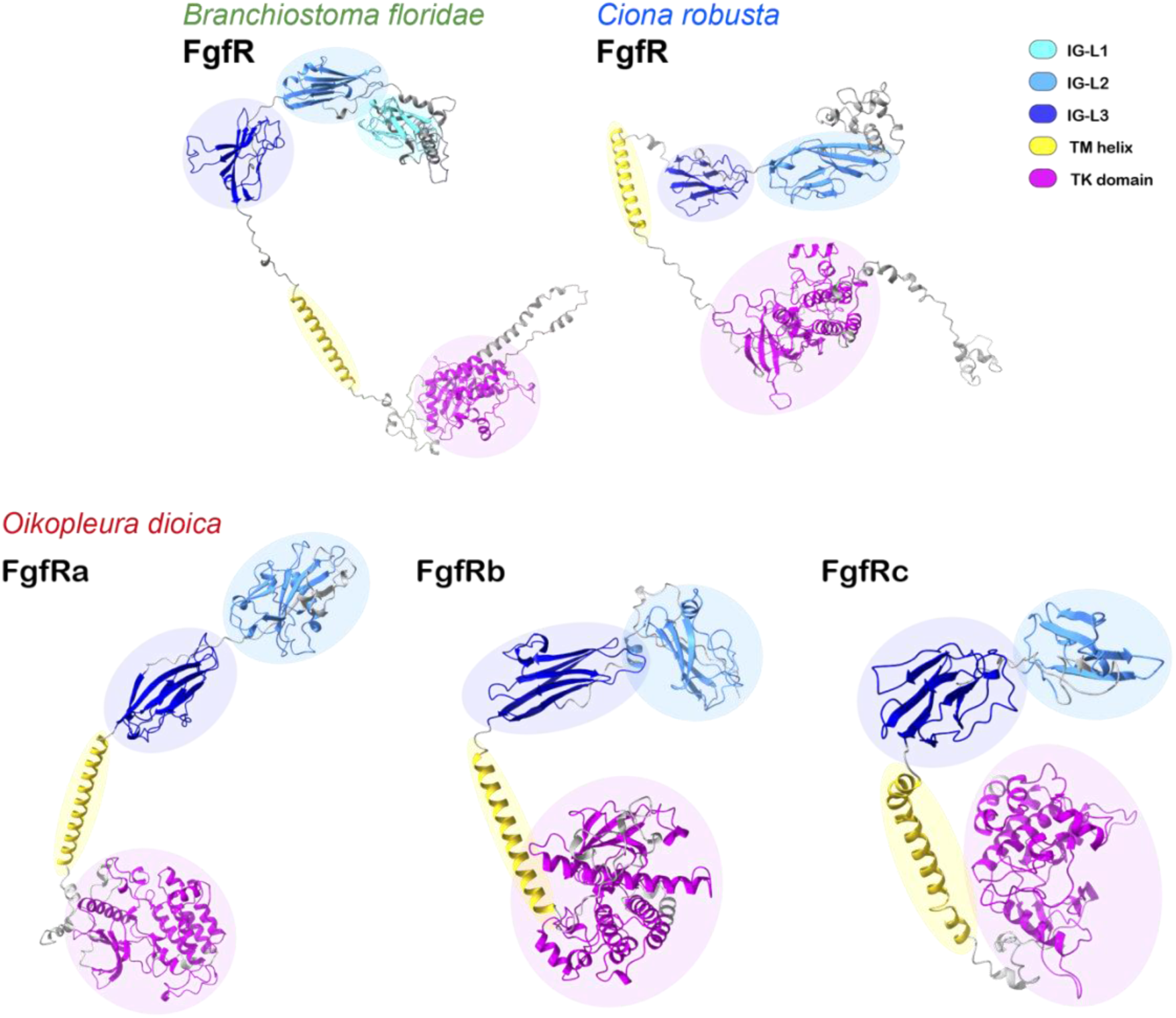
Three-dimensional models of the Fgf receptors. O. dioica FgfR models are based on the protein sequences of the annotations derived from this project. *B. floridae* FgfR model is based on the protein sequence in XP_035673320.1. *C. robusta* FgfR model is based on the protein sequence in NP_001037820.1. Models were manually coloured to mark the different domains and motifs that build each protein. An alternative colouring according to the local confidence score of the predictions is depicted in **Supplementary** Figure 1.

Regarding the intracellular portion that included the tyrosine kinase (TK) domain, the amino acid sequence was well conserved among the three *O. dioica* paralogs, although lower than that typically expected for a catalytic domain in other species (*O. dioica* FgfR TK domains exhibited ∼50% sequence similarity among paralogs, in contrast to 80-90% among the *H. sapiens* FgfR paralogs) (**Supplementary Table 2**). Additionally, the intron code was completely different among the three *O.dioica* FgfR paralogs (**Figure 3A**). This was most notable considering that the intron code was widely conserved among TK domain-containing protein orthologs, often used to classify a given TK domain-containing protein into a specific protein family (**Figure 3B**) (D’Aniello et al. 2008). In the extracellular regions that mediate ligand binding, *O. dioica* FgfRs showed strikingly low sequence similarity, with only 27–36% identity among paralogs, far below the 64–74% similarity observed among *H. sapiens* FgfR paralogs (**Supplementary Table 2**). When we compared the extracellular portion of human or *O. dioica* FgfRs with that of other chordates, we observed that *O. dioica* FgfRs in all cases displayed a lower similarity than human FgfRs, even with their closely related ascidian ortholog, highlighting the exceptionally rapid evolution of this domain in appendicularians (**Supplementary Table 3**).

**Figure 3.**
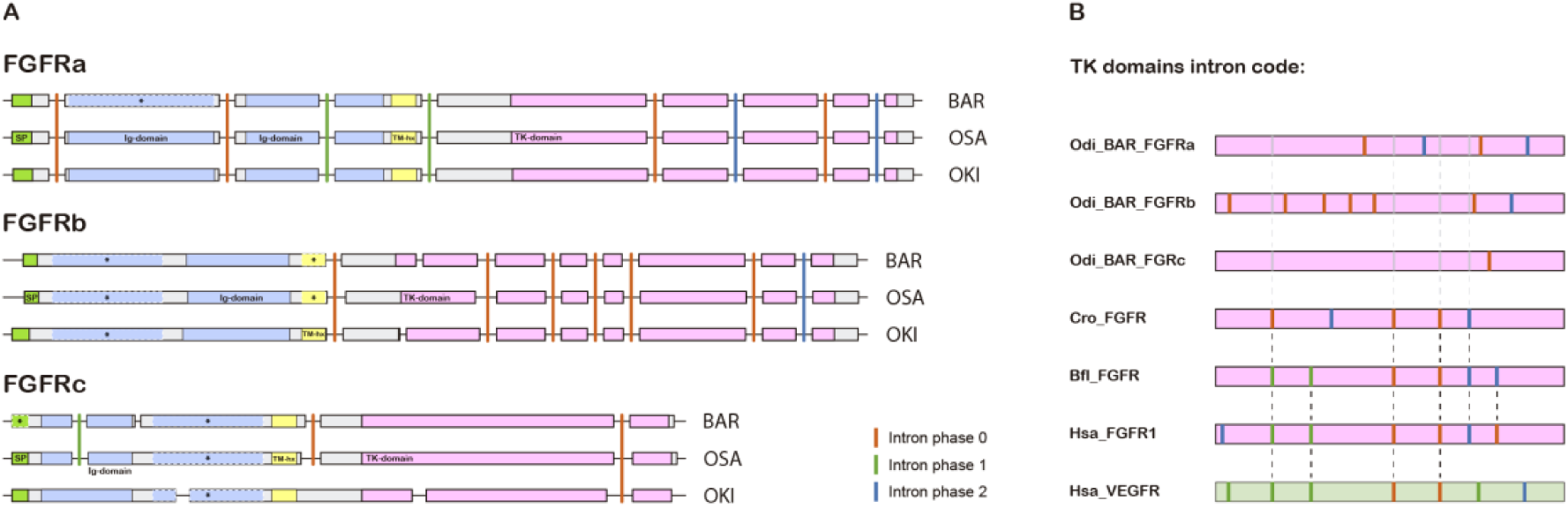
*O. dioica* FgfR gene structures. (A) Compared structures of FgfR orthologs from the three O. dioica cryptic species. Exons are scaled and depicted as boxes over the non-coding intronic regions (not scaled) depicted as lines. Vertical lines show the relative position of conserved introns, and their phase is indicated with a colour code. Coloured boxes with black solid lines over the grey background illustrate the conserved domains as identified by the InterProScan software. Coloured boxes with white dashed lines marked with an asterisk illustrate domains that were not identified by the InterProScan software, due to a remarked sequence divergency, and whose presence was inferred based on the proteins’ 3D structure prediction. (B) Intron code of *O. dioica* FgfR TK domains compared to other chordate orthologs. Solid dashed lines mark the posi:on of conserved introns, and shaded dashed lines mark the position of lost introns in *O. dioica* FgfRs TK domains.

This divergence was also evident among FgfR orthologs from the three cryptic *O. dioica* species. Similarity in the extracellular region was markedly reduced compared with the intracellular tyrosine kinase domain. BAR–OSA orthologs retained 90–95% identity, comparable to human–mouse FgfR comparisons, whereas BAR–OKI orthologs dropped to 79–83%, a range similar to human–chicken orthologs (70–90%) (**Supplementary Table 3**). Considering that humans and chickens diverged ∼310 million years ago (Khamsi 2004), while the cryptic *O. dioica* species split only ∼25 million years ago (Plessy et al. 2024), this pattern underscored the extraordinary evolutionary rate of *O. dioica*. This extreme divergence mirrored the high sequence variability observed in *O. dioica* Fgf9/16/20 ligands. Although further work is needed to determine ligand–receptor specificity, it is plausible that each of the three FgfRs has specialized to interact with one of the three major ligand clades: the basal Fgf9/16/20a family, the intronless signal-peptide-bearing Fgf9/16/20bcf group, or the intronless non–signal-peptide Fgf9/16/20de group (Sánchez-Serna et al. 2025).

### Distinct expression patterns of *O. dioica FgfR* paralogs through development suggest functional diversification

To understand the potential functional consequences of the expansion and diversification of the FgfR genes in *O. dioica*, we performed *in situ* hybridization assays to detect their expression throughout embryonic and larval development (**Figure 4**). *FgfRa* and *FgfRb* transcripts were strongly detected as early as in the oocytes, revealing that these two genes were part of the maternal component. For the *FgfRc* gene, on the other hand, we did not detect a clear expression signal until the 64-cell stage, and the embryos required long periods of incubation for the staining to be detected. These observations were consistent with the Gene Expression Matrix data available in the OikoBase, in which *FgfRa* (GSOIDG00008404001) and *FgfRb* (GSOIDG00009004001) displayed high levels of expression, especially in the earliest stages of development, while *FgfRc* (GSOIDG00010261001) expression was not detected (**Supplementary Table 4**) (Danks et al. 2013).

**Figure 4.**
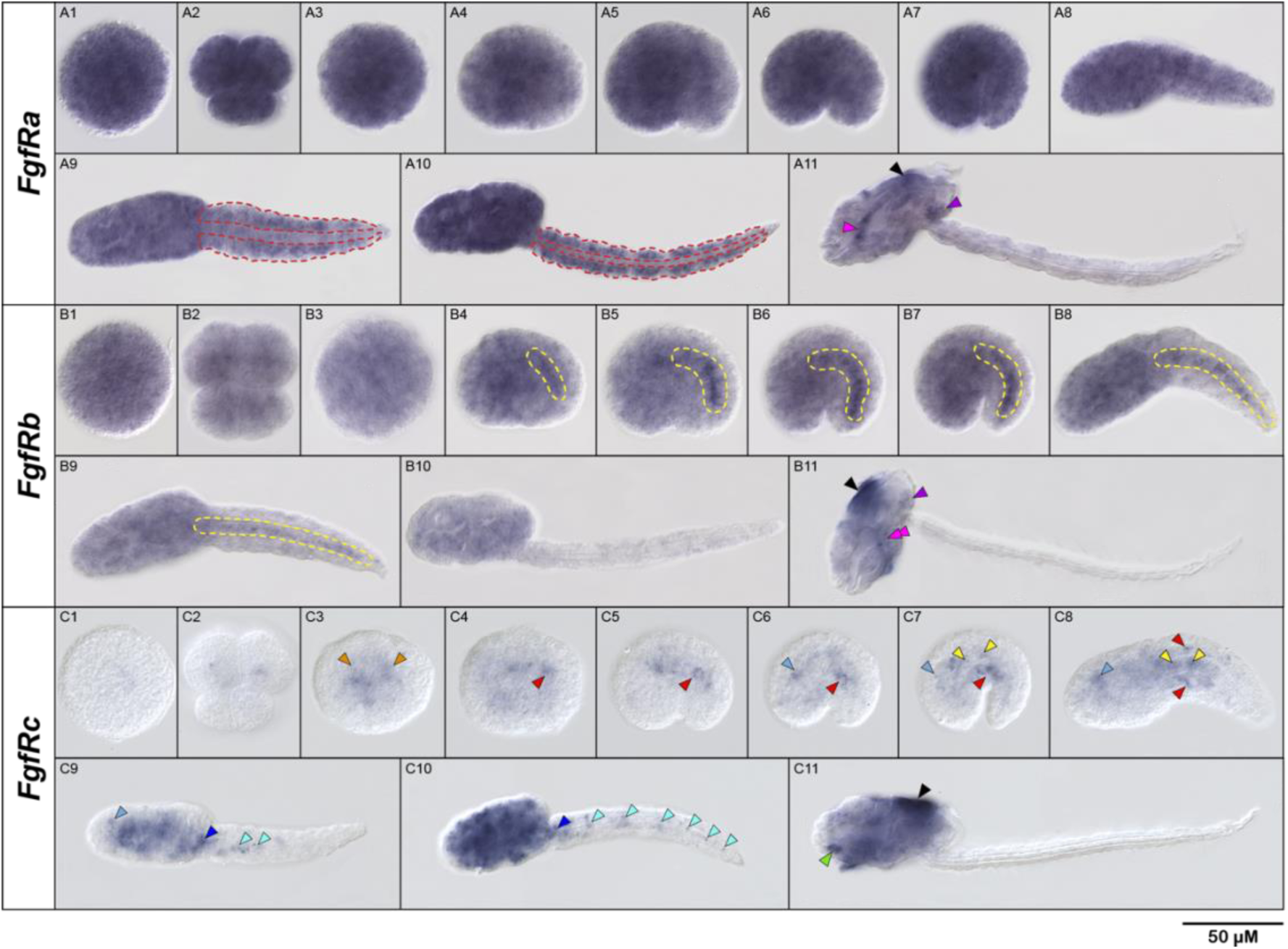
FgfR gene expression throughout *O. dioica* development. ABC1 are eggs, ABC2 are 8-cell embryos, ABC3 32-cell embryos, ABC4,5,6 and 7 are incipient, early, mid, and late tailbud embryos, respectively; ABC8,9,10 and 11 are just-, early-, mid- and late-hatched larvae, respectively. Images of tailbud embryos and larvae are left lateral views, with the anterior to the left and the dorsal to the top. Red dashed lines mark the muscles of the tail, yellow dashed lines mark the notochord, orange arrowheads mark endomesodermal precursor blastomeres, red arrowheads mark muscle cells, light blue arrowheads mark the developing brain, dark blue arrowheads mark the developing caudal ganglion, cyan arrowheads mark neuronal bodies in the nerve chord, yellow arrowheads mark notochord cells, magenta arrowheads mark the walls of the pharynx, magenta double arrowheads mark the pharyngeal slits, green arrowheads mark the ventral organ, purple arrowheads mark the gonad, and black arrowheads mark the oikoplastic epithelium.

The expression signal of *FgfRa* and *FgfRb* was ubiquitous in virtually all stages, although some obvious tissue specific domains with higher signal could also be distinguished. *FgfRa* expression signal was detected homogeneously until the early hatch stage, when a specific and more intense staining appeared in the muscle cells of the tail (**Figure 4 A9-10,** red dashed lines). This specific staining was maintained until the late hatch stage, the time at which the tail muscles become functional for the larvae to start swimming. Similarly, *FgfRb* expression signal was homogeneous in the embryos until the incipient tailbud stage, when a specific and more intense expression became evident in the developing notochord until the early hatch stage (**Figure 4 B4-9**, yellow dashed lines). This time frame coincided with some of the main processes of organogenesis of the notochord, including the proliferation, convergence and extension, fusion, and vacuolization of the notochord cells (Søviknes and Glover 2008). In contrast, *FgfRc* showed a restricted pattern of expression in all developmental stages examined. Its expression first became clear at the 64-cell stage, when the staining could be detected in some of the inner cells that give rise to endomesodermal derivatives (**Figure 4 C3**, orange arrowheads). In tailbud stages and just-hatched larvae, the *FgfRc* gene was asymmetrically expressed in the anteriormost left muscle cells in the tail (**Figure 4 C4-C8**, red arrowheads), and in the anterior region of the notochord (**Figure 4 C7-C8**, yellow arrowheads). *FgfRc* expression was also detected in the developing neural system, including the developing brain, the caudal ganglion, and neuronal bodies along the nerve cord from the mid-tailbud embryo to the mid-hatched larvae, when the expression signal became generalized in the trunk (**Figure 4 C6-9**, different tones of blue arrowheads)

In the late-hatched larvae, when most organs and structures were completing their development to become functional, the three *FgfR* genes showed novel specific expression domains. *FgfRa* and *FgfRb* expression signal was detected in a domain compatible with the gonad primordium (**Figure 4 A11 and B11**, purple arrowheads); *FgfRa* expression was also detected in the walls of the pharynx (**Figure 4 A11**, green arrowhead); and *FgfRc* expression was detected in the ventral organ (**Figure 4 C11**, green arrowhead). Moreover, at this late hatched stage all three *FgfR* genes showed a clear signal of expression in some fields the oikoplastic epithelium (**Figure 4 A11, B11 and C11**, black arrowheads), as did most of the *Fgf* genes (Sánchez-Serna et al. 2025).

Our results, therefore, showed that the three *FgfR* paralogs in *O. dioica* were expressed during embryonic and larval development, and the fact that they were expressed with different intensities in different structures suggested that they acquired different developmental functions after their expansion. While *FgfRa* and *FgfRb* expression signal was ubiquitous and their putative specific functions seemed to be mostly related to certain mesodermal structures (i.e. the tail muscles and the notochord), and endemesodermal derivatives (i.e. pharynx, ventral organ and gonad primordium), *FgfRc* expression seemed to become more specifically related to ectoneural derivatives (i.e. nervous system and the oikoblastic epithelium).

### Conservation and reduction of components of the transduction pathways

Upon activation of the FgfR by an Fgf ligand, a diverse array of proteins is recruited to the territory of the receptors TK domains and phosphorylated to propagate the signal through one or multiple transduction cascades. In this study, we asked whether the genetic rearrangements that shaped the diversity of *O. dioica Fgf* and *FgfR* genes were paralleled by changes in the intracellular components of the signaling system. We focused on the three major FgfR-associated pathways: the RAS/MAPK pathway, the PLCγ/PKC pathway, and the PI3K/AKT pathway (Goetz and Mohammadi 2013).

A careful genome survey revealed that most of the core components of the three abovementioned pathways were conserved in the genome of *O. dioica*, with the notable exception of the RAS/MAPK pathway (**Supplementary Table 4**) In the PI3K/AKT pathway, we identified orthologs of PI3KCA, PDK, and AKT; in the PLCγ/PKC pathway, orthologs of PLCγ and representatives of all four vertebrate PKC groups were present (Newton 2010). For the RAS/MAPK pathway, orthologs of Raf, MEK1/2, and ERK1/2 were identified, but none of the classical Ras proteins (H-Ras, K-Ras, or N-Ras) were detected in the genomes of any of the three cryptic *O. dioica* species (**Supplementary Table 4**).

The absence of classical Ras genes in *O. dioica* mirrored previous observations in ascidians, where M-Ras may substitute for classical Ras proteins during FgfR-mediated signaling in neural development (Keduka et al. 2009). To test for a similar mechanism, we searched for orthologs of M-Ras and other Ras family members. While no clear orthologs of M-Ras or R-Ras were identified, orthologs of Rheb, Ral, and Rap were present, along with additional Ras-like genes of uncertain assignment. Given the diversity of the Ras family (Wennerberg et al. 2005), and the precedent of functional substitution in ascidians, it is likely that *O. dioica* employed an alternative Ras member to mediate MAPK signaling. Thus, the absence of classical Ras did not necessarily imply disruption of the cascade, a view further supported by the conservation of its three downstream kinases (RAF, MEK1/2, and ERK1/2). *O. dioica* genomes also lacked genes encoding signal-transducing adaptor proteins. These proteins typically act as accessories to the main proteins in transduction cascades, facilitating protein-protein interactions to form larger signaling complexes (Luo and Hahn 2015). For instance, we observed the absence of *Fibroblast-growth-factor-receptor-substrate-2α* (*FRS2α*) and *Grb2-associated binder-1* (*Gab1*), the latter of which was also absent in ascidian *C. robusta*.

Altogether, our genome survey revealed that the main components of the three classical transduction pathways upon FgfR activation were present in *O. dioica*, which pointed to a conservation in the potential outcomes following activation of the receptors. The absence of classical *Ras* genes, as well as the absence of some of the adaptor proteins, suggested that the arrangement of these initial transduction complexes and the mechanisms by which the activation of the FgfR was transmitted to downstream kinases differ between *O. dioica* and vertebrates, but did not point to significant rearrangements on their putative functionalities.

### Developmental expression of the main components of RTK transduction pathways

To investigate potential relations between the Fgf/FgfR system and the downstream classical transduction pathways, we analyzed the expression pattern of some of their representative components throughout *O. dioica* development (**Figure 5**).

**Figure 5.**
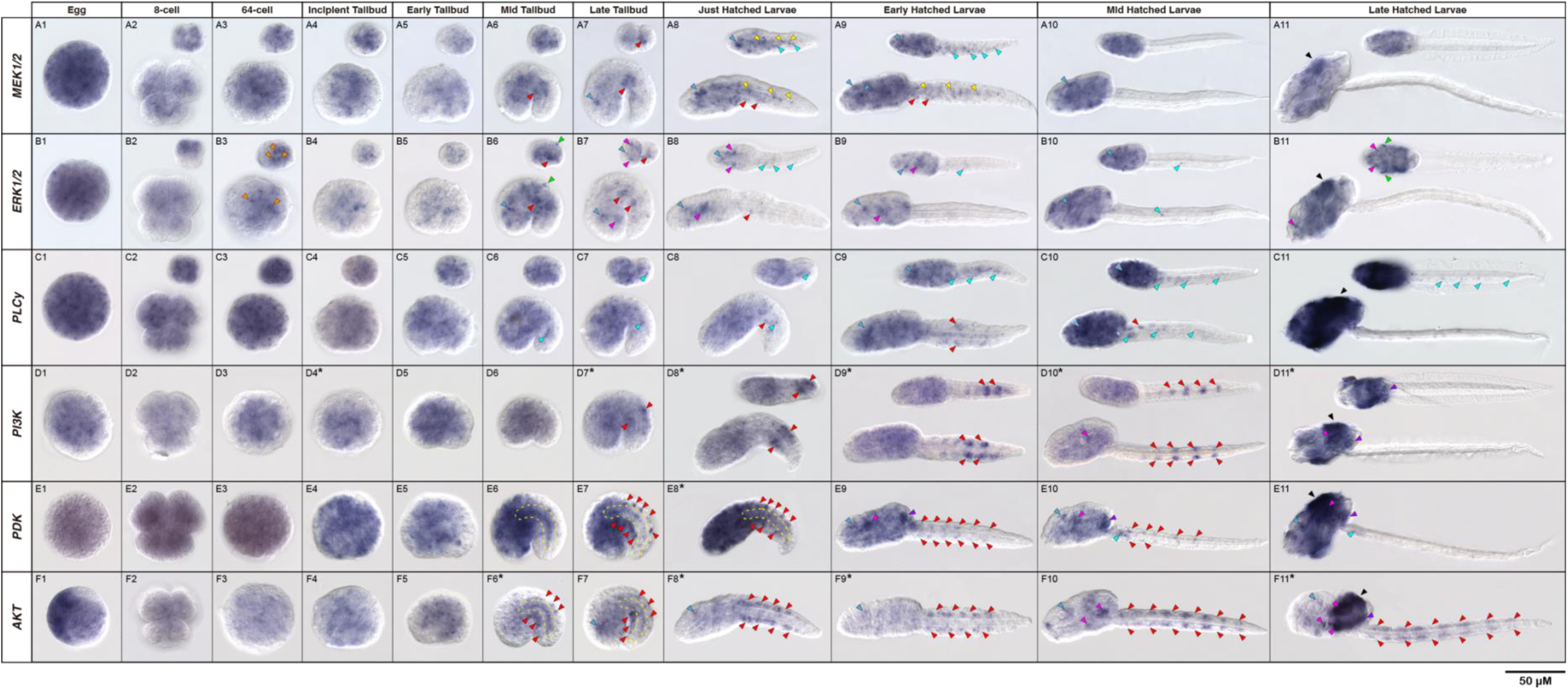
Developmental expression patterns of *O. dioica* genes involved in intracellular transduc0on pathways. (A-F) Expression panels of the genes selected as representatives of the three main pathways. Orange arrowheads mark endomesodermal precursor blastomeres, red arrowheads mark muscle precursors and muscle cells, light blue arrowheads mark the developing brain, cyan arrowheads mark neuronal bodies in the caudal ganglion and the nerve chord, yellow arrowheads mark notochord cells and yellow dashed lines mark the whole notochord, magenta arrowheads mark undetermined domains in the trunk, green arrowheads mark epithelial domains, and purple arrowheads mark the gonad primordium. All panels except those marked with an asterisk are left lateral views of the embryo or larvae with the anterior to the left and the dorsal to the top. Pannels marked with an asterisk are right views of the embryo or larvae, mirrored for aesthetic purposes. All insets are dorsal views of the same embryo pictured in the panel.

#### a) The MAPK pathway

In the MAPK pathway, *O. dioica* MEK1/2 and ERK1/2 transcripts were strongly detected from egg to tailbud stage, suggesting a maternal origin and ubiquitous early distribution (**Figure 5 A1–5, B1–5**). ERK1/2 also showed zygotic expression in vegetal blastomeres at the 32-cell stage, marking endomesodermal precursors (**Figure 5 B3**). From the tailbud stage onwards, expression weakened and became restricted to the trunk and anterior tail (**Figure 5 A6–9, B6–10**). Specific domains included the developing brain (**Figure 5 A7–9, B6–10**, blue arrowheads), anterior ventral tail muscles (**Figure 5 A6–9, B6–10**, red arrowheads), the notochord (**Figure 5 A8–9**, yellow arrowheads), and neuronal bodies along the nerve cord (**Figure 5 A8–9, B6–10**, cyan arrowheads). In addition, ERK1/2 was expressed in presumptive pharyngeal slit regions from the late tailbud to early hatchling stage (**Figure 5 B7–9**, magenta arrowheads) and in a unique pair of epidermal tail cells in late tailbud embryos (**Figure 5 B6**, green arrowhead).

#### b) The PLCy pathway

For the PLCγ/PKC pathway, PLCγ expression was detected ubiquitously from the egg to the mid-tailbud stage (**Figure 5 C1–6**). During these stages, embryos stained rapidly and homogeneously, with no clear evidence of specific expression domains. From the late tailbud to early hatchling stages, expression became more restricted to the trunk and anterior tail, with stronger signals in the developing brain, nerve cord, and tail muscle cells (**Figure 5 C7–9**, blue, cyan, and red arrowheads). In mid- to late-hatchlings, the trunk retained a generally uniform signal with only the brain standing out against the background. In contrast, the tail no longer showed generalized expression, but specific PLCγ expression was detected in neuronal bodies along the nerve cord (**Figure 5 C10–11**, cyan arrowheads).

#### c) The PI3K pathway

In the PI3K/AKT pathway, *PI3KCA*, *PDK*, and *AKT* were maternally expressed and ubiquitous until tailbud stages, after which more restricted and tissue-specific expression pattern were revealed. In these later stages, *PI3KCA* showed stronger staining in the gut and gonad primordium (**Figure 5 D10–11**), PDK in the brain, gut, and gonad primordium (**Figure 5 E9–11**), and *AKT* in the brain, posterior endoderm, stomach lobes, posterior pharynx floor, migrating buccal glands, and gonad primordium (**Figure 5 F7–11**). Besides these expression patterns in the trunk, the *PI3KCA*, *PDK* and *AKT* genes appeared to be specifically expressed in the tail muscle cells in all hatching stages; but while *AKT* and *PDK* seemed to be expressed indistinctly in all the muscle cells, *PI3K* showed a dynamic expression pattern with different expression intensities in the cells along the tail. This pattern suggested that the PI3K/AKT pathway may play a specific role in axial cell identity and tail muscle development.

Altogether, these patterns indicated that FgfR downstream pathways were largely maternally provided, with later tissue-specific refinement, and suggested roles in brain, trunk, and tail muscle development as well as pharyngeal and epidermal patterning.

### Attenuators of the FGF signaling: evolution of the Sprouty/SPRED genes in appendicularians

The fact that the main components of transduction pathways displayed such a generalized expression pattern, especially in the earlier stages of development, encouraged us to search for downstream genes that could serve as markers for the activation of the Fgf receptors. Two of such potential genes were *Sprouty* and *SPRED* (*Sprouty-related Proteins with an EVH1 Domain*), which encode non-catalytic attenuators of RTK signaling in vertebrates (Kim and Bar-Sagi 2004; Neben et al. 2019). Sprouty and SPRED are closely evolutionarily related, as their proteins share an exclusive cysteine-rich domain, the SPRY domain, close to their C-terminus. While Sprouty proteins do not carry any other conserved domain, SPRED proteins also bear an EVH1 domain close to their N-terminus (Kawazoe and Taniguchi 2019). This EVH1 domain is closely related to that of Ena/VASP proteins, a conserved family of actin regulatory proteins (Krause et al. 2003). Thus, SPRED proteins look like a chimera of an Ena/VASP and a Sprouty protein. Despite these two related proteins have been generally associated to a “switching” mechanism by negative feedback-regulation of RAS-MAPK activation, their molecular mechanisms and other potential functions are not fully understood, and Sprouty proteins have been found to also regulate the PLCγ response and other transduction pathways apart from the MAPK cascade (Nutt et al. 2001; Sivak et al. 2005; Ayada et al. 2009; Chow et al. 2009; Akbulut et al. 2010).

Our searches in the genome of *O. dioica* allowed us to identify a putative ortholog for *Sprouty*, but we could not find any clear ortholog for *SPRED*. To test the possible loss of *SPRED*, we searched for all the orthologs for *Sprouty*, *SPRED*, and *Ena/VASP* genes in appendicularians, ascidians, vertebrates and cephalochordates, and performed a phylogenetic reconstruction with all the sequences retrieved (**Figure 6A**). Our results confirmed the lack of *SPRED* in all appendicularians species surveyed, as none of the identified sequences containing either a SPRY or a EVH1 domain were grouped with SPRED proteins. As this gene was present in all other chordates, our results indicated a loss of the *SPRED* gene in the appendicularians lineage after their split from the rest of tunicates and prior to their radiation (**Figure 6B**).

**Figure 6.**
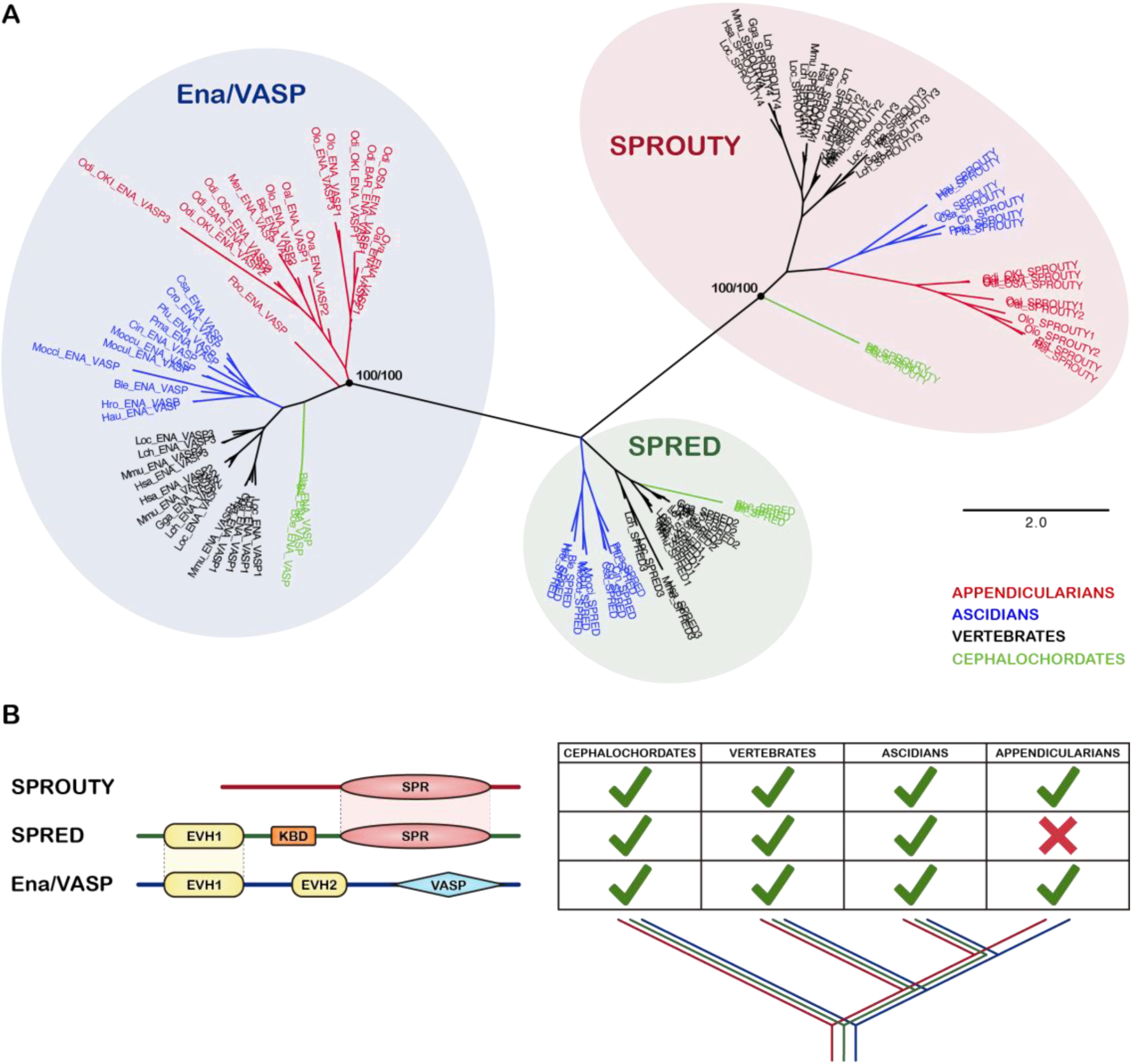
Evolution of Sprouty and Spred proteins in appendicularians. (A) Maximum likelihood phylogenetic inference of Ena/VASP, Sprouty and Spred proteins from cephalochordates (green), vertebrates (black), ascidians (blue) and appendicularians (red). Node support values correspond to SH-aLRT support (%) / ultrafast bootstrap support (%). The scale bar indicates amino-acid substitutions. (B) Graphic representation of Sprouty, Spred, and Ena/VASP domain architectures and their presence in the chordate clades surveyed. The loss of Spred proteins only affects the appendicularian lineage.

### Developmental expression of *Sprouty* in *O. dioica*

We next analyzed the expression of the *Sprouty* gene throughout development in *O. dioica* **(Figure 7A)**. From the egg to the 32-cell stage embryo we did not find any signal of gene expression, which indicated that it was not part of the maternal mRNA contribution. At the 32-cell stage, when gastrulation began, the expression signal started to be detected in the nuclei of three pairs of blastomeres. In order to determine more confidently their identity, we performed a one-colour double WMISH assay using *Brachyury* as a marker of the notochord precursors, whose positions were precisely determined in the 32-cell and 64-cell embryos **(Figure 7B)**. The expression signal revealed that *Sprouty* started to be expressed in three territories in the 32-cell embryo, according to Delsman’s/Conklin’s nomenclature (Stach et al. 2008): the B22/B6.1 pair of vegetal blastomeres, that will give rise to endomesodermal derivatives including the whole endodermal strand; the A11/A6.4 and A12/A6.3 pairs within the neural plate, that will develop into most of the nervous system; and the a11/a6.8 pair of animal blastomeres that will give rise to part of the epidermis. At the 64-cell stage, once the embryos were undergoing gastrulation, *Sprouty* expression persisted in the whole neural plate, in the nuclei of endomesodermal precursor cells derived from the B22/B6.1 pair, and in the animal blastomeres derived from the a11/a6.8 pair. Moreover, two new expression domains appeared: one in the b7.9 and b7.10 pairs of blastomeres, that at this time were already restricted to the muscle fate, and the other in a posterior pair of animal blastomeres that will develop into epidermis **(Figure 7 A3-4 and Figure 7B)**.

**Figure 7.**
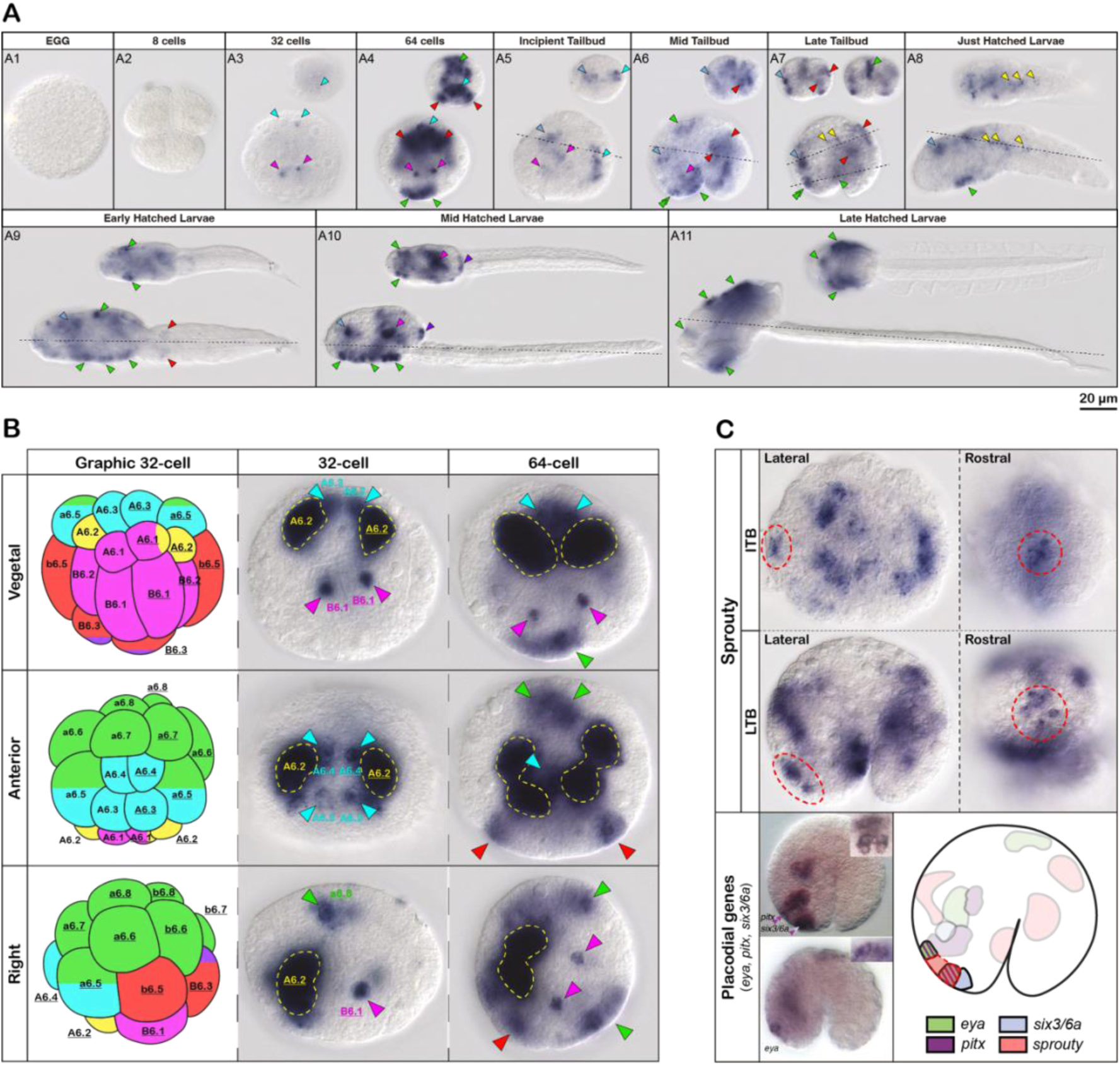
Analyses of Sprouty gene expression. (A) Sprouty gene expression throughout O. dioica development. All panels are left lateral views of the animals with the anterior to the left and the dorsal to the top. All insets are dorsal views of the animals with the anterior to the left at the focal plane indicated with dashed lines in the main panel. The scale bar does not account for the insets. Cyan arrowheads mark the neural plate and the developing nerve chord, magenta arrowheads mark endomesodermal precursors and their derivatives, red arrowheads mark muscle precursors and muscle cells, light blue arrowheads mark the developing brain, green arrowheads mark epidermal domains, green double arrowheads mark the oral placode, yellow arrowheads mark notochord cells, and purple arrowheads mark the gonad primordium. (B) Left column is a graphic representation of the 32-cells embryo displaying blastomeres with the same colour code as the arrowheads in (A) and numbered according to Conklin’s nomenclature. Mid and right columns are 32-cell and 64-cell embryos, respectively, outcoming from a one-colour double WMISH for Sprouty and Brachyury genes. Yellow dashed lines mark the notochord precursor cells expressing Brachyury, and coloured arrowheads mark Sprouty expression domains. Each row displays a different orientation of the embryos, as indicated in the lettering. (C) Upper panels are detailed images of Sprouty gene expression in the region of the oral placode, surrounded by a red dashed line. In the bottom, images of WMISH for placodal genes (i.e. eya, pitx and six3/6a) were extracted from Bassham & Postlethwait, 2005. Bottom right panel is a graphic representation of a late tailbud embryo in lateral left view with expression domains for eya, pitx, six3/6a and Sprouty collapsed.

In later stages, the expression signal of *Sprouty* was detected mainly in structures derived from the blastomeres that were stained in the 32-cell and 64-cell embryo. These were neural structures, including the posterior region of the developing neural cord **(Figure 7 A5, cyan arrowheads)** and the brain during most of its development **(Figure 7 A5-10, blue arrowheads)**; muscle cells of the tail, especially the three anteriormost pairs in the mid and late tailbud stages **(Figure 7 A6-7, red arrowheads)**; undetermined endomesodermal structures within the trunk **(Figure 7 A5-10, magenta arrowheads)**; and different regions of the trunk epidermis, including the oikoplast in the late hatched larvae **(Figure 7 A5-11, green arrowheads)**. All these domains overlapped, at least partially, with expression domains of *Fgf* and *FgfR* genes, what supported that Sprouty might be involved in the cellular response to the activation of the FgfR. Moreover, *Sprouty* expression signal could also be observed in the anterior most region of the notochord in the late tailbud and just hatched larvae stages **(Figure 7 A7-8, yellow arrowheads)**, similarly to the specific staining observed for several Fgf9/16/20 paralogs (Sánchez-Serna et al. 2025).

Among the epidermal regions stained, a specific rostral domain in the mid and late tailbud embryos was revealed as of special interest **(Figure 7 A6-7, double green arrowheads)**. This epidermal region is known to develop into the oral placode, whose homology to vertebrate placodes was confirmed by the expression of placodal markers such as *Eya*, *Pitx* and *Six3/6* (Bassham & Postlethwait, 2005). The clear expression of *Sprouty* in some cells within this region suggested that its function was compatible with that described in vertebrates as an attenuator of the RTK pathway, and could therefore be essential for the proper patterning and development of placodes **(Figure 7C)**.

## DISCUSSION

Our analyses on the evolution of the FgfR gene and the downstream transduction pathways, together with our previous work on FGF ligands (Sánchez-Serna et al. 2025), reveal that the evolution of the FGF pathway in appendicularians follows two contrasting trajectories: while the signaling components, namely Fgf ligands and their receptors, have undergone a remarkable remodeling including expansion and diversification, the intracellular transduction machinery has remained largely conserved, and in several cases simplified. This dichotomy reflects fundamental differences in the evolvability of the two functional layers that constitute any signaling pathway: (1) the extracellular signaling interface, responsible for implementing specific developmental instructions through ligand–receptor interactions, and (2) the intracellular transduction core, which integrates and propagates these signals using deeply conserved molecular modules (Pires-daSilva and Sommer 2003; Schaefer et al. 2014; Grandchamp and Monget 2018). *Oikopleura dioica* provides a clear example of how these two layers evolve under distinct constraints.

### Lineage-specific duplication and divergence of FgfRs parallels the evolution of Fgf ligands

We find that the three FgfR paralogs of *O. dioica* originate from appendicularian-specific duplications and retain canonical receptor organization despite remarkable sequence divergence. Their evolutionary history mirrors the previously described expansion of the two surviving Fgf ligand subfamilies, which suggests diversification of ligand–receptor pairs (Sánchez-Serna et al. 2025). The parallel FgfR amplification to the Ffg ligand expansion in appendicularians, therefore, follows the same pattern as the one observed in vertebrates after the Fgf and FgfR expansion associated with the two-rounds of whole-genome duplication, in sharp contrast with the single-copy condition of FgfR and Fgf gene families in all other non-vertebrate chordates (Itoh and Ornitz 2004; Oulion et al. 2012).

Sequence and intron–exon comparisons further illustrate how rapidly appendicularian Fgf receptors have evolved. Their extracellular domains show low similarity to those of other chordates and even among paralogs, and their intron structures are highly divergent. However, the overall receptor layout remains comparable to that of ascidians, and the intracellular tyrosine kinase domain is considerably more conserved than the extracellular portion. This pattern of innovation in the ligand-binding regions combined with strong conservation of the intracellular core supports the idea that receptors act as flexible evolutionary interfaces: their outer domains can change to create new signaling specificities, while the essential features needed for intracellular activation remain preserved (Pires-daSilva and Sommer 2003; Grassot et al. 2006; Brunet et al. 2016).

### Diversification of developmental expression supports functional innovation

Expression profiling indicates spatial and temporal subfunctionalization among *O. dioica* FgfRs. The maternally supplied and broadly expressed paralogs, *FgfRa* and *FgfRb*, contrast with *FgfRc*, which is strictly zygotic and confined to specific neural, mesodermal, and epithelial territories. These complementary patterns point to a redistribution of ancestral receptor functions: FgfRa and FgfRb likely support early embryogenesis, whereas FgfRc appears to have acquired more specialized roles later in development. Beyond their maternal expression, FgfRa and FgfRb also display distinct late-stage expression, FgfRa in muscle cells and FgfRb in the notochord, as well as in specific organ domains in late hatchlings.

Notably, several FgfR expression territories coincide with tissues that have undergone lineage-specific innovation in appendicularians, such as the oikoplastic epithelium and the pharyngeal slits. This correspondence reinforces a broader evo-devo principle: diversification of ligand–receptor repertoires often accompanies morphological novelty, enabling developmental programs to be rewired through changes in signaling machinery (Kirschner and Gerhart 1998; Babonis and Martindale 2017). The association between receptor expression and these lineage-specific traits provides a basis for future studies on how the FgfR diversification may have contributed to the rewiring of developmental programs underlying morphological innovations in appendicularians.

### Intracellular transduction components are highly conserved or even simplified

While the signaling machinery has been notably remodeled in comparison to other chordates, the intracellular transduction machinery has remained remarkably stable. The principal effectors of the PI3K/AKT, PLCγ/PKC, and RAS/MAPK pathways are retained in *O. dioica* and show broadly conserved developmental expression. However, two reductions stand out: the absence of classical Ras proteins and the loss of several adaptor molecules. These changes suggest a trend toward simplification rather than expansion, consistent with patterns previously described for *O. dioica* (Seo et al. 2004; Denoeud et al. 2010; Albalat and Cañestro 2016; Ferrández-Roldán et al. 2021).

This contrast aligns with a well-established evolutionary principle: intracellular transduction modules experience stronger purifying selection than ligand–receptor components. Kinases, adaptors, and other core effectors carry out fundamental biochemical functions conserved across eukaryotes, from yeast to animals; alterations in these components typically perturb basic cellular physiology rather than developmental patterning (Widmann et al. 1999; Pires-daSilva and Sommer 2003). Consequently, their evolutionary change is limited, and functional innovation tends to accumulate instead at the signaling interface.

### Two evolutionary trends within a single pathway: signaling vs. transduction

The contrasting behavior of the two layers of the FGF pathway offers direct evidence for differential evolvability between signaling functions and transduction functions. Signaling functions (ligands and receptors) evolve rapidly, generating developmental diversity. Their interactions are modular and relatively easily rewired, allowing organisms to innovate at the level of tissue communication without altering core cellular physiology. By contrast, transduction functions (intracellular cascades) evolve slowly due to their pleiotropy and the central role they play in cellular homeostasis.

In *O. dioica* and other appendicularians, this contrast is especially pronounced. Fgf ligands and receptors have expanded and diverged despite pervasive genome reduction, whereas intracellular RTK cascades have retained their core organization, experiencing only limited gene loss that does not disrupt the essential architecture of the pathway. In vertebrates, these contrasting trends are often obscured by the two rounds of whole-genome duplication (2R-WGD), which expanded all components simultaneously. As a result, the differential evolvability of signaling vs. transduction layers is masked by the global duplication of both modules. The fast-evolving appendicularians, which exhibit both the retention of ancestral chordate traits and the emergence of lineage-specific innovations, and which have evolved without genome-wide duplication events, represent an excellent model system for investigating these evolutionary dynamics.

## ACKNOWLEDGEMENTS

We thank all present and past team members on CC’s laboratory for assistance and fruitful discussions, specially to Sebastian Artime Paoletti for running the Oikopleura facility in the University of Barcelona. We thank to Centres Científics i Tecnològics de la UB for sea water supply and sequencing services.

## AUTHOR CONTRIBUTIONS

Conceptualization: CC, GS; formal analysis: GS; funding acquisition: CC; investigation: GS, PB, AF, MF, AAB, LRL, NPT; project administration and supervision: CC; writing: GS, CC.

## SUPPLEMENTARY FILES

**Supplementary Figure 1. Three-dimensional models of the Fgf receptors. *O. dioica* FgfR models are based on the protein sequences of the annotations derived from this project.** B. floridae *FgfR* model is based on the protein sequence in XP_035673320.1. C. robusta *FgfR* model is based on the protein sequence in NP_001037820.1. Models are coloured according to their local QMEANDisCo score along the protein sequence. Coloured shades in the background mark the different domains and motifs that build each protein according to the colour code in the legend.

**Supplementary Table 1. Genes cloned in this project.** Primers used, length of the insert, DNA used as a template, and genome used for the design of the primers.

**Supplementary Table 2. Compared FgfR identity and similarity between *H. sapiens* paralogs and *O. dioica* paralogs.** Percentages were inferred separately for the intracellular TK domain and the extracellular portion of the FgfRs.

**Supplementary Table 3. Interspecies FgfR identity and similarity.** Percentages were inferred separately for the intracellular TK domain (only *O. dioica* cryptic species) and the extracellular portion of the FgfRs. Species included: *Oikopleura dioica* (Odi), *Homo sapiens* (Hsa), *Ciona robusta* (Cro), *Branchiostoma floridae* (Bfl), *Mus musculus* (Mmu), *Gallus gallus* (Gga) and *Lepisosteus oculatus* (Loc).

**Supplementary Table 4. Gene expression matrix values for *FgfR* genes.** Extracted from the OikoBase (hIp://oikoarrays.biology.uiowa.edu) (Danks et al., 2013). Expression units are not specified in the source.

**Supplementary Table 5. Search of genes involved in intracellular transduction pathways.** The human gene was first used to identify the ortholog in *C. robusta*, and subsequently human and ascidian genes were used as queries to search the ortholog in *O. dioica*. Hyphens indicate genes not found through the RBBH method.

