## Supplementary figures and images for "Downstream effects of the “Less, but More” Fgf signaling in *Oikopleura dioica*: Fgf receptor expansion and RTK pathway simplification"

### SuppFig_1_FgfR-3Dmodels.png

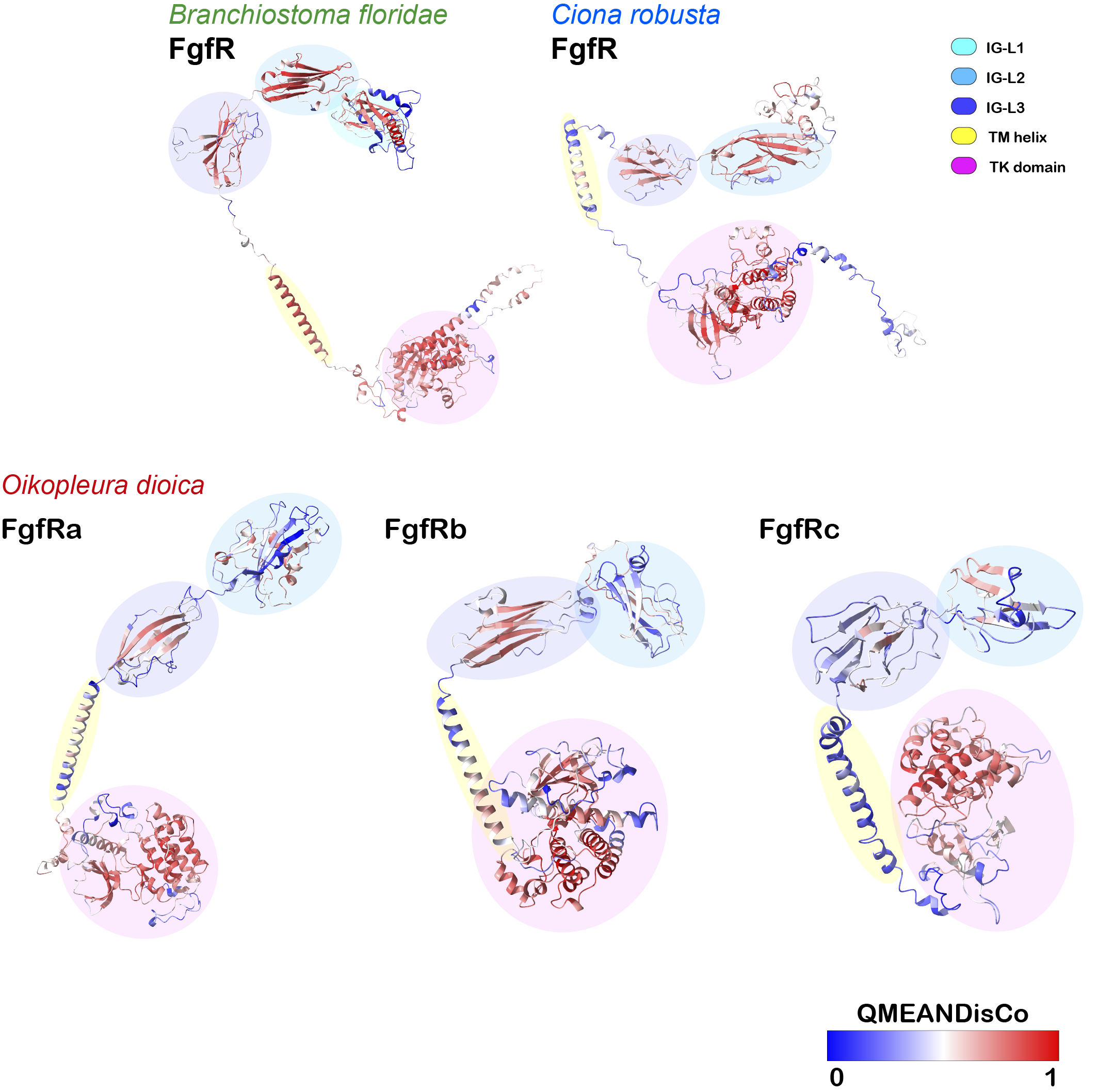
